# RNA sequencing on muscle biopsy from a 5-week bedrest study reveals the effect of exercise and potential interactions with dorsal root ganglion neurons

**DOI:** 10.1101/2021.08.11.455963

**Authors:** Amelia J. McFarland, Pradipta R. Ray, Salman Bhai, Benjamin Levine, Theodore J. Price

## Abstract

Lack of physical activity is a predictor of poor health outcomes that can be prevented or reversed by exercise. Sedentary lifestyle, chronic disease or microgravity can cause muscle deconditioning that then has an impact on other physiological systems. An example is the nervous system, which is adversely affected by decreased physical activity resulting in increased incidence of neurological problems such as chronic pain. We sought to better understand how this might occur by conducting RNA sequencing experiments on muscle biopsies from human volunteers in a 5-week bed-rest study with an exercise intervention arm. We also used a computational method for examining ligand-receptor interactions between muscle and human dorsal root ganglion (DRG) neurons, the latter of which play a key role in nociception and are generators of signals responsible for chronic pain. We identified 1352 differentially expressed genes (DEGs) in bed rest subjects without an exercise intervention but only 132 DEGs in subjects with the intervention. Thirty-six genes were shared between the exercise and no intervention groups. Among 591 upregulated muscle genes in the no intervention arm, 26 of these were ligands that have receptors that are expressed by human DRG neurons. We detected a specific splice variant of one of these ligands, placental growth factor (PGF), in deconditioned muscle that binds to neuropilin 1, a receptor that is highly expressed in DRG neurons and known to promote neuropathic pain. We conclude that exercise intervention protects muscle from deconditioning transcriptomic changes, and prevents changes in expression of ligands that might sensitize DRG neurons that promote pain. Our work creates a set of actionable hypotheses to better understand how deconditioned muscle may influence the function of sensory neurons that innervate the entire body.

## Introduction

Skeletal muscle is highly adaptive and undergoes cellular physiology and phenotype changes as well as transcriptomic reprogramming, in particular in response to extended reductions in physical activity (Booth et al.; Patterson et al., 2018). Accordingly, in situations whereby mechanical stimuli and load are significantly reduced – such as prolonged bedrest or immobilization, exposure to microgravity environments, or a highly sedentary lifestyle – deconditioning responses are observed (Baldwin, 1996; Brocca et al., 2012; Booth et al., 2017). Inactivity and reduced physical activity have been associated with an array of maladaptive responses, with sedentary time considered an important, independent contributor to the development of metabolic and cardiovascular disease, as well as all-cause mortality (Wilmot et al., 2012; Dempsey et al., 2016; Gabriel and Zierath, 2019). On the other hand, exercise has clear health benefits, which include cardiovascular and metabolic systems and extend to neurological outcomes, such as pain relief (Sluka et al., 2018) and protection from development of chronic pain (Grace et al., 2016; Leung et al., 2016).

Following reductions in loading, skeletal muscle experiences loss of mass and fiber size, muscle fibers shift towards a fast phenotype, myofibril content decreases, reduced synthesis of muscle protein occurs, and muscle protein is degraded by a proteasomal mechanism (DeMartino and Ordway, 1998; Adams et al., 2003; Gallagher et al., 2005; Haddad et al., 2005; Hastings et al., 2012; Krainski et al., 2014b; Phillips and McGlory, 2014). Previous studies have demonstrated that high-intensity exercise countermeasures, such as rowing ergometry and resistive strength exercises, are capable of preserving cardiac and skeletal muscle mass and limit cardiovascular deconditioning associated with prolonged bed rest (Hastings et al., 2012; Krainski et al., 2014b). Indeed, the integrated cardiovascular benefits associated with rowing ergometry mean that it has been suggested as a strategy to counter severe deconditioning scenarios, such as prolonged spaceflight (Hastings et al., 2012; Krainski et al., 2014b), or chronic diseases such as the Postural Orthostatic Tachycardia Syndrome (Fu et al., 2010).

While the vast majority of studies in this field to date have focused on functional and/or structural changes associated with bedrest or limb immobilization, gene expression profiling tools have also permitted exploration of the transcriptional changes which accompany muscle deconditioning, and how exercise may lessen or counteract these (Pillon et al., 2020). Microarray technology has been used in both acute limb immobilization as well as acute and chronic bed rest studies to explore associated transcriptional changes (Urso et al., 2006; Abadi et al., 2009; Chopard et al., 2009; Lammers et al., 2012; Fernandez-Gonzalo et al., 2020), while more comprehensive RNA-sequencing / RNA-seq (which allows for a deeper investigation of the transcriptome compared to microarray analysis) has only been employed in short term bed rest studies (≤ 14 days) (Mahmassani et al., 2019a; Mahmassani et al., 2021). Despite the link between physical inactivity and onset of musculoskeletal complications and systemic symptoms, knowledge of the mechanisms that underlie this link are not well understood. Moreover, it is not clear whether deconditioned muscle may contribute to disease mechanisms beyond the cardiovascular or musculoskeletal systems, for instance through an action on neurological systems (Booth et al., 2017), which could contribute to the somatic hypervigilance that is present in deconditioning syndromes (Cutsforth-Gregory and Sandroni, 2019). We have recently developed computational methods that allow for detailed examination of pharmacological interactions between any tissue and human dorsal root ganglion (DRG) neurons (Wangzhou et al., 2021). These neurons play a critical role in pain signaling and tactile and thermal sensation, and signals derived from muscle may interact with these neurons to mediate neurological effects of exercise or deconditioning.

The purpose of the present study was to use RNA sequencing to gain mechanistic insight into transcriptional changes which occur following muscle deconditioning in long-lasting bed rest. We analyzed muscle biopsy samples from healthy individuals which had been previously collected as part of a 5-week head-down-tilt bed rest investigation (Krainski et al., 2014b). Using a discovery-based approach, we performed bulk RNA-seq on muscle samples, and sought to identify genes, pathways and processes which were differentially expressed between individuals who underwent complete bed rest, and those who performed a high-intensity exercise routine during their bed rest. Additionally, we evaluated the potential impact of transcriptional changes in muscle on DRG neurons using a genome-wide ligand-receptor pair database curated for pharmacological interactions relevant to neuro-immune systems (Wangzhou et al., 2021). This work represents the longest duration of human physical inactivity analyzed via RNA sequencing technology, and integrates this new knowledge into the context of potential impacts on DRG neurons.

## Methods

### Subjects

Skeletal muscle biopsies were collected at the Institute for Exercise and Environmental Medicine. Methods and subject characteristics for this study have been previously described (Krainski et al., 2014b), shown in Table 1 and in Supplementary Data, Sheet 1. Subjects were healthy, nonsmoking, adults aged 20-54 years. Subjects were randomly assigned to one of two groups: bed rest only (BR-CON; n=9, 1 female), or bed rest with exercise countermeasure (BR-EX; n=16, 2 female). All subjects underwent-6-degree head down tilt bed rest for 35 consecutive days. BR-EX subjects received an exercise countermeasure consisting of rowing ergometer training on 6 days/week and biweekly resistance training for the duration of the bed rest period. A needle muscle biopsy of 300 mg tissue on average was obtained from the midbelly of the right (pre) and left (post) vastus lateralis muscle before and after the five week bed rest period (Krainski et al., 2014b). Part of each biopsy was used for a previously published study (Krainski et al., 2014b); at least 50 mg per sample remained for use in the present study.

**Table 1:**
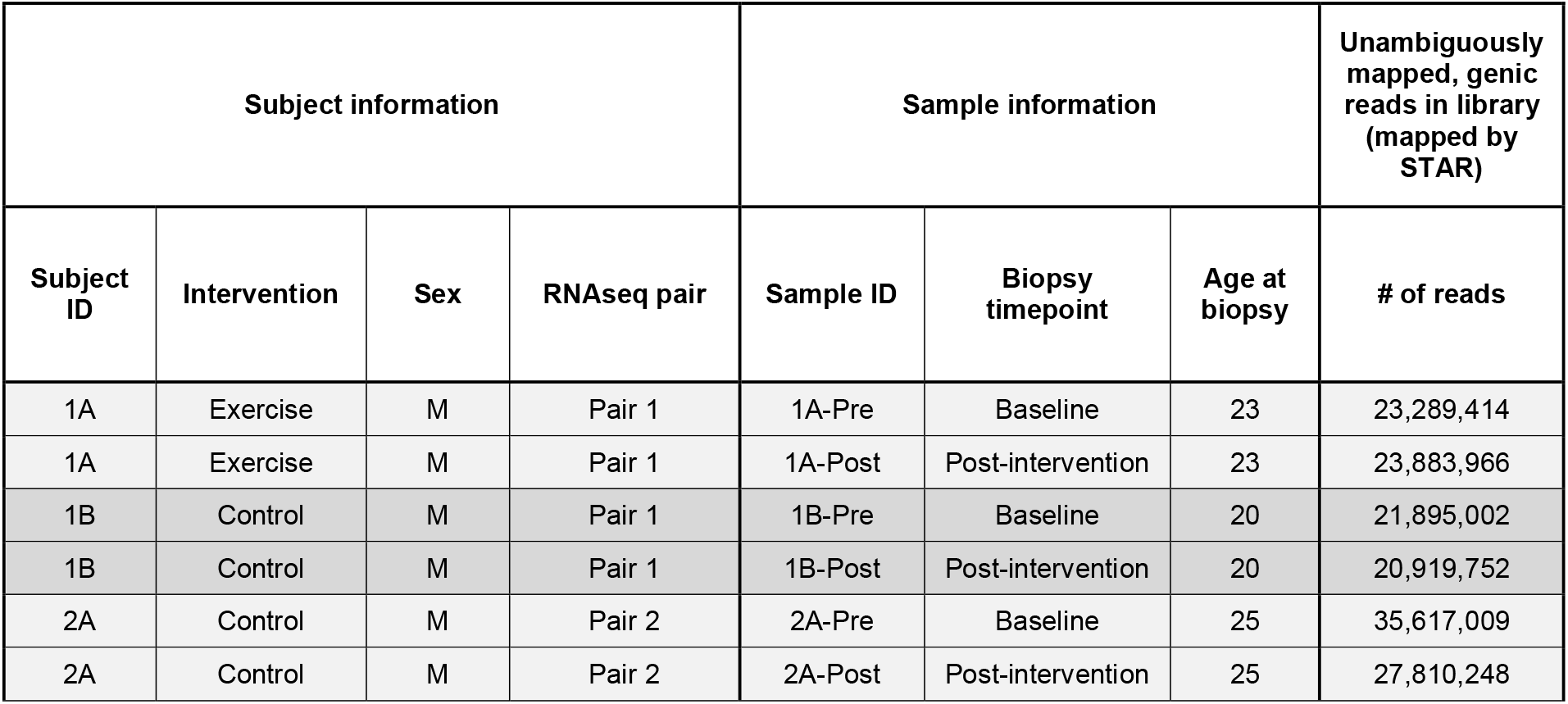
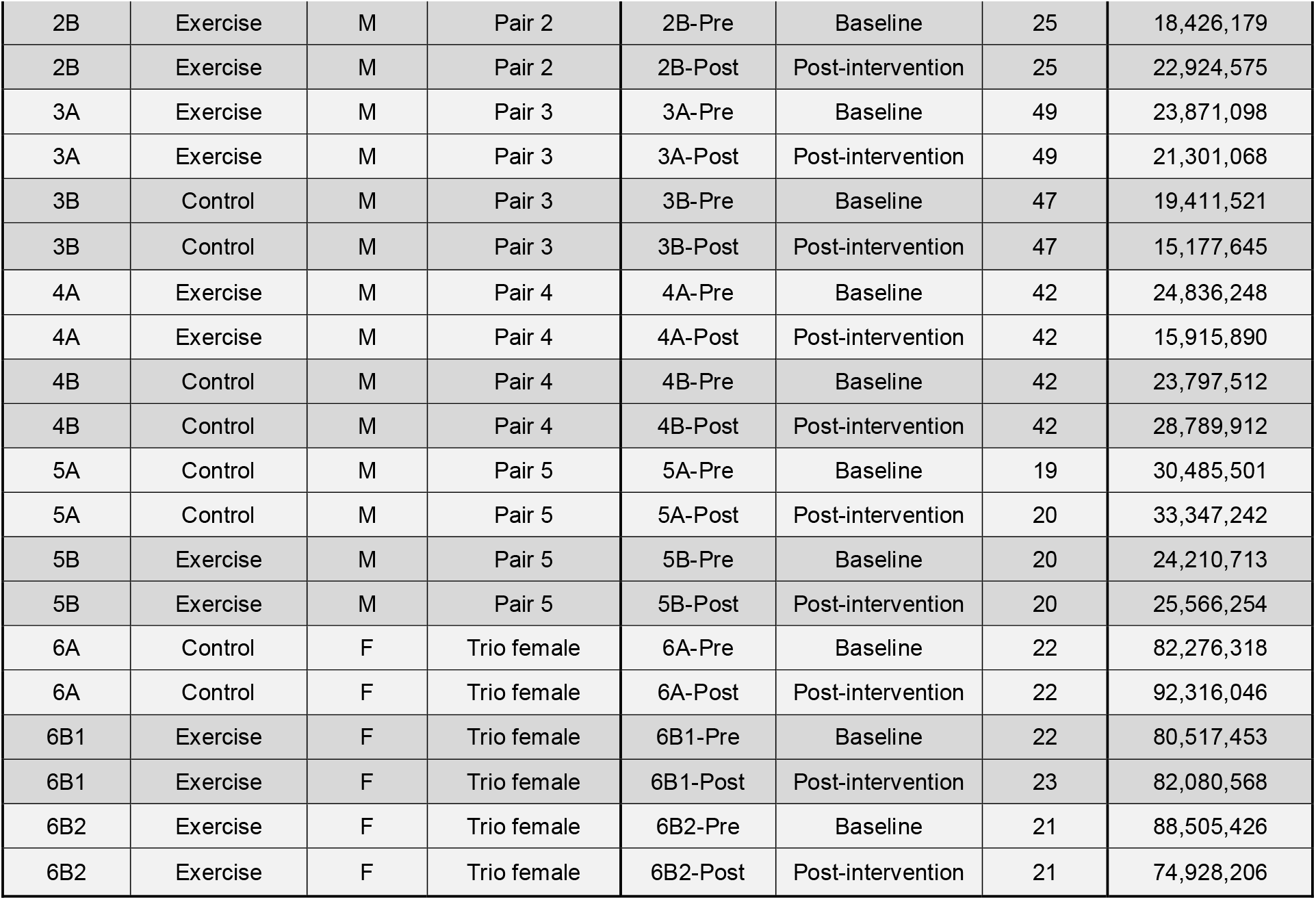
Sample sheet for all biopsies from subjects included in this study

### Library Generation and Sequencing

Based on available muscle following initial ultrastructural and histochemical analysis (Krainski et al., 2014a) biopsy samples from 13 subjects underwent RNA sequencing, consisting of all three female samples (1 BR-CON, 2 BR-EX), and ten male samples (5 BR-CON, 5 BR-EX; matched for age and exercise tolerance at baseline). Researchers performing sequencing and subsequent analyses were informed of subject pair matches and whether samples were from baseline or post-intervention, but were blinded to intervention received by individual subjects (i.e. the researchers did not know whether subjects were sedentary or exercised during bedrest until the first round of analysis was complete). Following RNA extraction (RNeasy Plus Universal Mini Kit, Qiagen), RNA yield was quantified using a Nanodrop system (ThermoFisher Scientific), and RNA quality assessed by fragment analyzer (Advanced Analytical Technologies). Stranded mRNA library preparation and sequencing were completed at the University of Texas at Dallas (UT Dallas), TX, USA. 76-bp, single-end sequencing of RNA-seq libraries was performed on the Illumina Hi-Seq sequencing platform.

### Mapping and Quantification of RNA-seq data

RNA-seq read files (fastq files) were checked for quality by FastQC (Babraham Bioinformatics) and read trimming was done based on Phred score and per-base nucleotide content, with trimmed, 60 bp reads from the 13 – 72 bp positions being used for downstream analysis. Trimmed reads were mapped to the reference transcriptome GENCODE v27 (Frankish et al., 2019), and its underlying reference genome GRCh38.p10 retaining only uniquely mapped reads by using the STAR (v2.7.3a) aligner (Dobin et al., 2013) with the following command line mapping parameters:

~~~
STAR --runThreadN 20 --runMode alignReads --genomeLoad NoSharedMemory --
outMultimapperOrder Random --outSAMtype BAM SortedByCoordinate --outSAMattributes All
--outSAMattrIHstart 0 --outSAMprimaryFlag AllBestScore --outBAMcompression-1 --
outBAMsortingThreadN 20 --outFilterType BySJout --outFilterMultimapNmax 10 --
outFilterMismatchNoverReadLmax 0.06 --twopassMode Basic --quantMode GeneCounts --
alignSJoverhangMin 5 --alignSJDBoverhangMin 3 --alignIntronMax 1000000 --
alignMatesGapMax 1000000 --outFilterIntronMotifs RemoveNoncanonical --
outSAMstrandField intronMotif --outFilterScoreMinOverLread 0.3 --
outFilterMatchNminOverLread 0.3 --readFilesCommand zcat --alignSoftClipAtReferenceEnds
No --alignEndsType EndToEnd
~~~

Relative abundance per sample (in Transcripts per Million or TPM) was calculated from the mapped reads using Stringtie (Pertea et al., 2015). Genes were filtered to only retain protein-coding genes and TPMs were re-normalized per sample to a million based only on the set of coding genes A fold-change ratio (post intervention / baseline) was calculated for each coding gene in every subject. (Supplementary Data, Sheet 2).

### Blinded analysis

The set of subjects were initially partitioned into pairs of subjects with matching sex and comparable ages (Table 1). For each subject pair, exactly one was BR-CON, while the other was BR-EX. Blinded researchers were told which sample belonged to which pair, but which member of the pair received which intervention was not revealed to researchers. The female samples were not included as part of the blinded analysis.

Hierarchical clustering of baseline and post-intervention samples was performed separately using TPMs of all protein coding genes, using (1-Pearson’s correlation coefficient) as the distance metric (Figures 1A and 1B). Multiple strategies were utilized to identify potential genes of interest from the blinded analysis of the dataset. Firstly, genes with the potential for differential regulation in the control and exercise groups were identified. This was achieved through identifying genes within each subject pair where one subject had a post intervention / baseline absolute log_2_ fold-change of ≥ 0.585 (corresponding to a fold change of ≥ 1.5 or ≤ 0.66), while the other subject had either no change (fold change between 0.66 and 1.5) or a fold change trend in the opposing direction. Genes that consistently appeared in the list for each subject pair were identified by intersecting the gene lists for all 5 subject pairs (Supplementary Data, Sheet 3). Additionally, genes with the potential for differential expression between baseline and post-intervention samples, irrespective of the intervention, were also identified. Lastly, genes with the potential for dysregulation due to bedrest were identified. This was achieved through calculating the relative variance (ratio of variance to mean) of TPMs values across sample pairs; the top 500 genes with the highest relative variance identified for each sample pair. Genes common to the top 500 highest relative variance genes in all sample pairs were also identified (Supplementary Data, Sheet 4). The relative variance and fold change gene lists generated through the blinded analysis were then compared, and genes that appeared in both lists were used to create a final gene set (Supplementary Data, Sheet 5).

**Figure 1:**
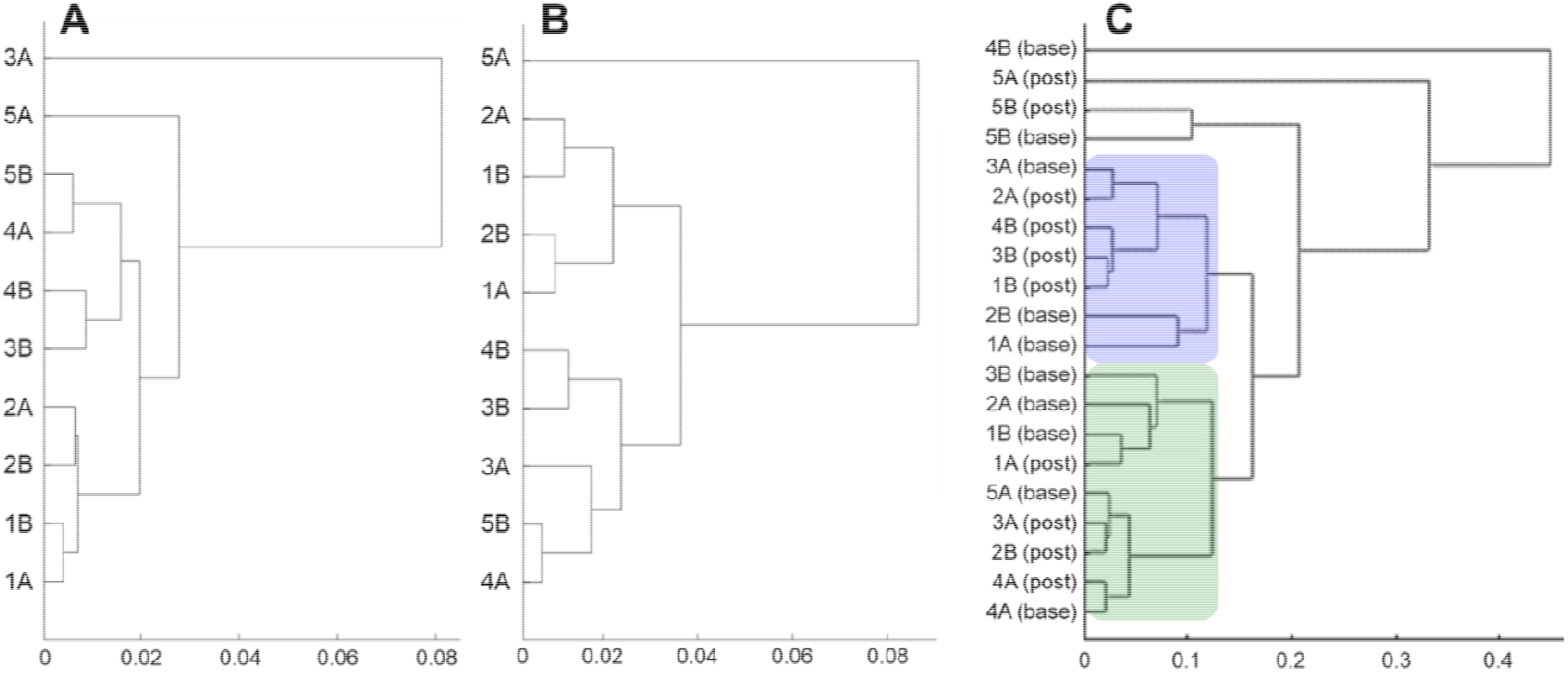
Dendrogram based on hierarchical clustering of TPMs from male subjects (n=10) with predicted groupings in blue and green. (A) All protein coding genes from paired subjects at baseline. (B) All protein coding genes from paired subjects post-intervention. (C) Protein coding genes meeting fold-change criteria (as identified from the blinded analysis) from paired subjects both at baseline and post-intervention. The distance (calculated as 1 – Pearson’s correlation coefficient between the vectors of coding gene TPMs in two samples) fitted to an ultrametric tree is shown on the x-axis.

To predict the intervention groupings for the male subjects, hierarchical cluster analysis of the TPMs for the fold-change gene list (Supplementary Data, Sheet 3; Figure 1C) in baseline and post-intervention samples together were used, and were also contrasted with previously described hierarchical cluster analysis of protein coding gene TPMs post-intervention (Figure 1B). Post-intervention samples for 4 subject pairs (pairs 1 – 4) were segregated into two clusters (Figure 1C), suggesting which subjects may have received the same intervention, Post-intervention samples for the 5^th^ pair (pair 5) were outliers in the clustering, and 5A and 5B were manually assigned to the two groups based on gene expression trends. Based on the predicted subject groupings, which group may have received which intervention was also predicted through comparison of this intersected gene set to previous bed rest datasets (Brocca et al., 2012; Mahmassani et al., 2019b).

Coding of subjects and samples (Table 1) was kept at the Levine lab while sequencing analysis was done in the Price lab. The Levine lab revealed the coding only after blinded data analysis from the Price lab was completed.

### Unblinded Differential Expression Analysis

After completion of the blinded analysis, researchers were unblinded to the interventions of each subject. Differentially expressed genes reported in the formal analysis in this report were identified using the R/Bioconductor package DEseq2, a well-established differential expression analysis tool which models read counts using a negative binomial distribution, and models dispersion as a function of the expression level (Love et al., 2014). Data were analyzed using a nested design to identify group-specific condition effects for BR-EX and BR-CON subjects post-intervention *vs*. baseline. Protein-coding genes with a TPM > 0.1, and an adjusted p-value of < 0.05 were used to identify differentially expressed gene sets for each condition (Supplementary Data, Sheets 6 and 7) P-value adjustment in DEseq2 is based on controlling the False Discovery Rate to <= 0.05 using the Benjamini-Hochberg correction, using the *p*.*adjust* function in limma / R (Wettenhall and Smyth, 2004). Gene set enrichment analysis on the differentially expressed gene (DEG) sets was performed using the Enrichr framework (Kuleshov et al., 2016). DEGs with potential for effects on DRG neurons were identified using a genome-wide ligand-receptor pair database curated for pharmacological interactions. An interactome was generated through intersecting the list of DEGs and their expression levels with a ligand-receptor pair list as previously described (Wangzhou et al., 2021).

### Identification of splice variants in PGF gene

Alternative splicing analysis was only performed for the exon retention / skipping event in PGF that caused gain / loss of heparin binding functionality in the encoded peptide respectively. Uniquely mapped bridge reads on the correct strand were counted using IGV (Thorvaldsdottir et al., 2013) for the relevant exon - exon junctions. Reads that overlapped the relevant exon that were identified this way were pooled for control and exercised subjects separately, and normalized to calculate the relative abundance of the alternatively spliced exon in question (in Reads per Kilobase per Million mapped reads). The number of mapped reads that were used for the normalization was the number of unambiguously mapped, genic reads that was mapped by the STAR aligner (Dobin et al., 2013) (relevant transcript and exon ids are provided in Supplementary Data, Sheet 1).

## Results

### Blinded analysis of bed rest RNA sequencing data

Of 58,763 total genes in the reference genome, we detected 36,072 genes in our samples. All samples yielded high quality raw sequence data. After filtering for protein-coding genes only, 17,730 unique genes remained.

Hierarchical cluster analysis of all male baseline samples showed high homology across all samples, and between most pairs (Figure 1A). Post-intervention, several non-paired subjects were seen to cluster together (Figure 1B), including subjects 4A and 5B, subjects 3B and 4B, subjects 1A and 2B, and subjects 2A and 1B. Through the use of our differential fold-change algorithm, where subjects within a pair demonstrated inconsistent or divergent gene expression post-intervention, we identified 263 genes which were consistently dissimilar between all subject pairs (Supplementary Data, Sheet 3). Additionally, we identified genes that were consistently trending in the same direction after bedrest irrespective of intervention (Supplementary Data, Sheet 3). Calculation of relative TPM variance across all subject pairs identified 141 genes which were consistently in the top 500 highest relative variance genes (Supplementary Data, Sheet 4). There were 17 genes which appeared consistently on all blinded analysis gene lists (Supplementary Data, Sheet 5).

Hierarchical cluster analysis of gene lists generated through fold change and relative variance computations similarly suggested high homology between most samples, particularly paired subjects, at baseline; however, post-intervention samples were observed to cluster primarily in non-paired groupings, consistent with previous hierarchical cluster analysis of all gene data. Based on the consistency of these post-intervention non-pair groupings, it was predicted that subjects 1B, 2A, 3B, 4B and 5A belonged to one group, while subjects 1A, 2B, 3A, 4A and 5B belonged to the other group. The pattern of expression for the 12 genes consistently expressed in all gene lists of interest were then reviewed for the two groups. Based on expression trends in previous bed rest studies (in particular, *XIRP1, TNFRSF12A*, and *HSPB1*) (Brocca et al., 2012; Mahmassani et al., 2019b), it was hypothesized that the control (bed rest only) group consisted of subjects 1A, 2B, 3A, 4A, and 5B, and that the intervention (bed rest plus exercise) group consisted of subjects 1B, 2A, 3B, 4B, and 5A. Following completion of the blinded analysis, investigators were unblinded to the subject grouping and intervention. The blinded analysis correctly predicted the grouping for all ten subjects, but incorrectly predicted the intervention received.

### Unblinded differential expression analysis of RNA sequencing data

Following unblinding of subject intervention, DESeq2 was used to determine which genes were differentially expressed between control and intervention groups for the male and female subjects. DESeq2 analysis found 1,352 differentially expressed genes (DEGs) in subjects in the BR-CON group (591 up-, 761 down-regulated post intervention vs. baseline, *p* < 0.05; see Supplementary Data, Sheet 6). In BR-EX subjects only 132 DEGs were identified (69 up-, 63 down-regulated post intervention vs. baseline, *p* < 0.05; see Supplementary Data, Sheet 7). Of the DEGs identified, 36 genes were consistent between the two groups; all other genes were differentially expressed in the BR-CON group were not significantly altered in subjects who received the exercise countermeasure. Twenty-two DEGs had consistent differential expression patterns in both BR-CON and BR-EX groups (16 up-regulated in both groups, 6 down-regulated in both groups), and an additional 14 genes had opposing trends between the groups (i.e., were up-regulated in one group, and down-regulated in the other). Of those genes which were differentially expressed in both BR-CON and BR-EX groups, *LBP* and *LRRC52* were both in the top-ten down-regulated genes when ranked by magnitude of fold change (Table 2), while *KIT* was found to be a top ten up-regulated gene for both groups.

**Table 2:**
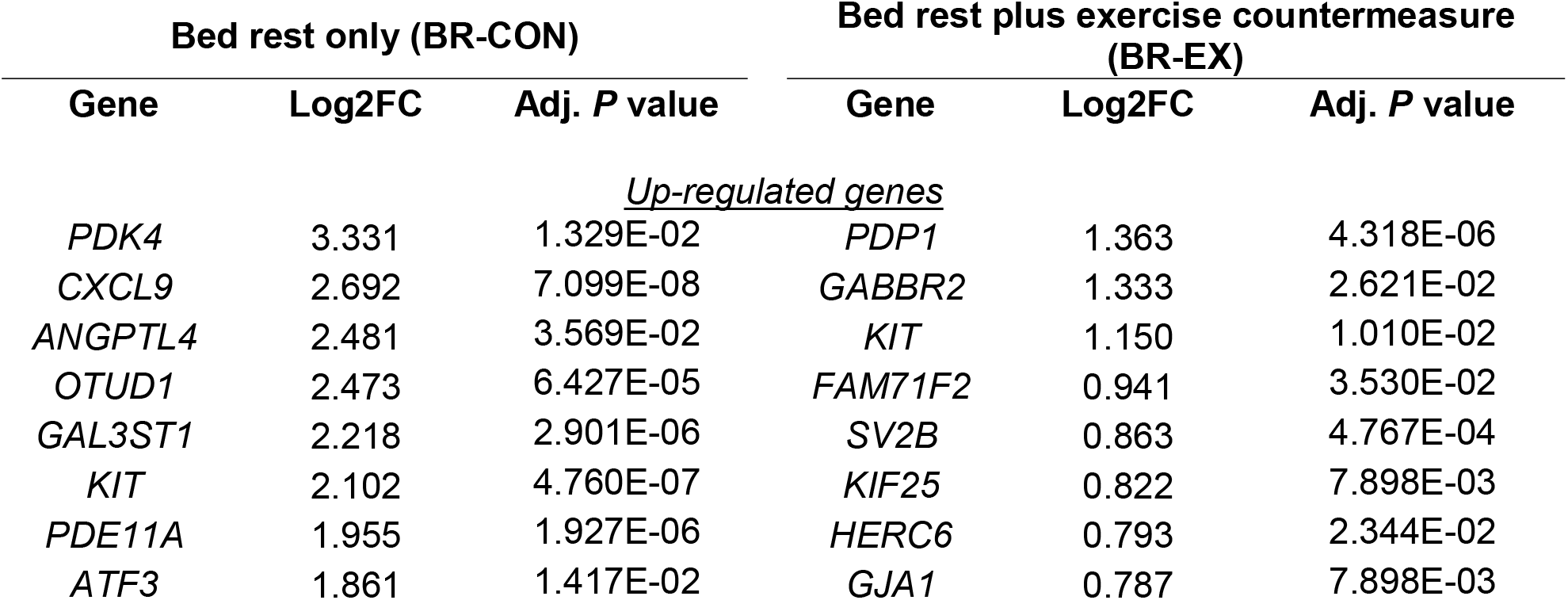
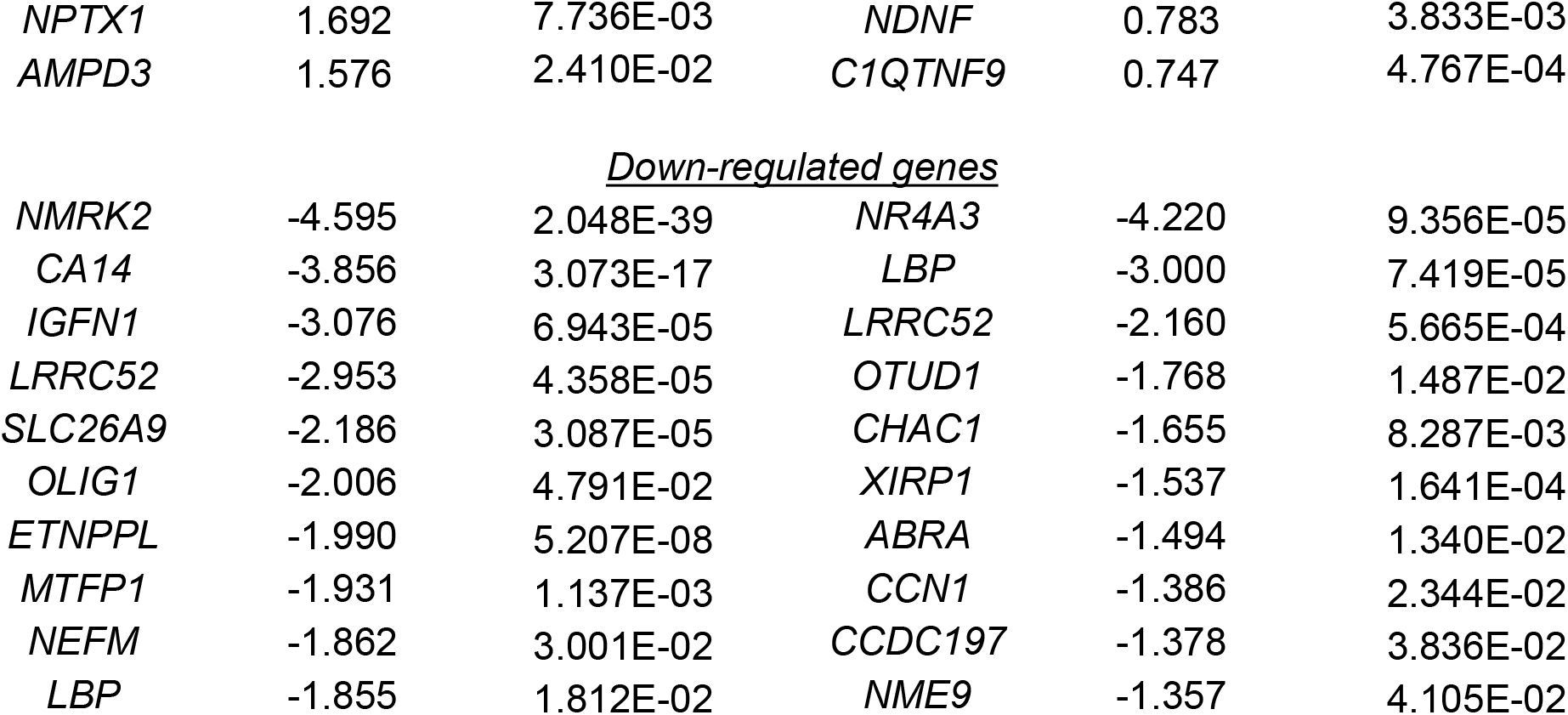
Top 10 genes up-regulated and down-regulated by 35 days of bed rest in BR-EX and BR-CON subjects. Genes are ordered by log_2_ of mean fold change value from baseline to post-intervention (Log2FC).

### Functional implications for gene expression change

The potential functional implications of gene expression changes were then explored through performing enrichment analysis on the DEGs from each group. In BR-CON, key up-regulated pathways identified by functional enrichment analysis included interferon-mediated signaling pathways, transcriptional regulation pathways, and antigen processing. Key pathways enriched by down-regulated DEGs in BR-CON related to mitochondrial respiration and electron transport (Table 3). In contrast, pathways enriched by down-regulated DEGs in BR-EX included transcriptional regulation pathways, cytokine-mediated signaling, and fatty acid oxidation regulation; pathways enriched by up-regulated DEGs included multiple pathways relating to endothelial cell development, proliferation and regulation (Table 4).

**Table 3:**
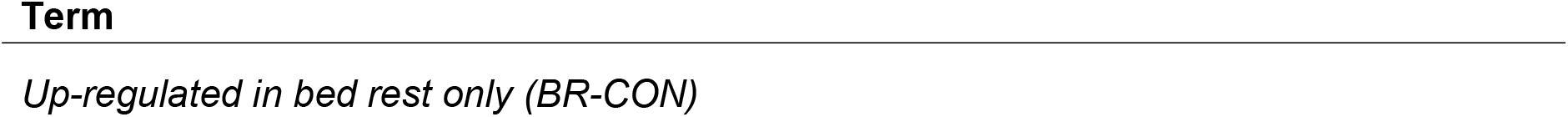
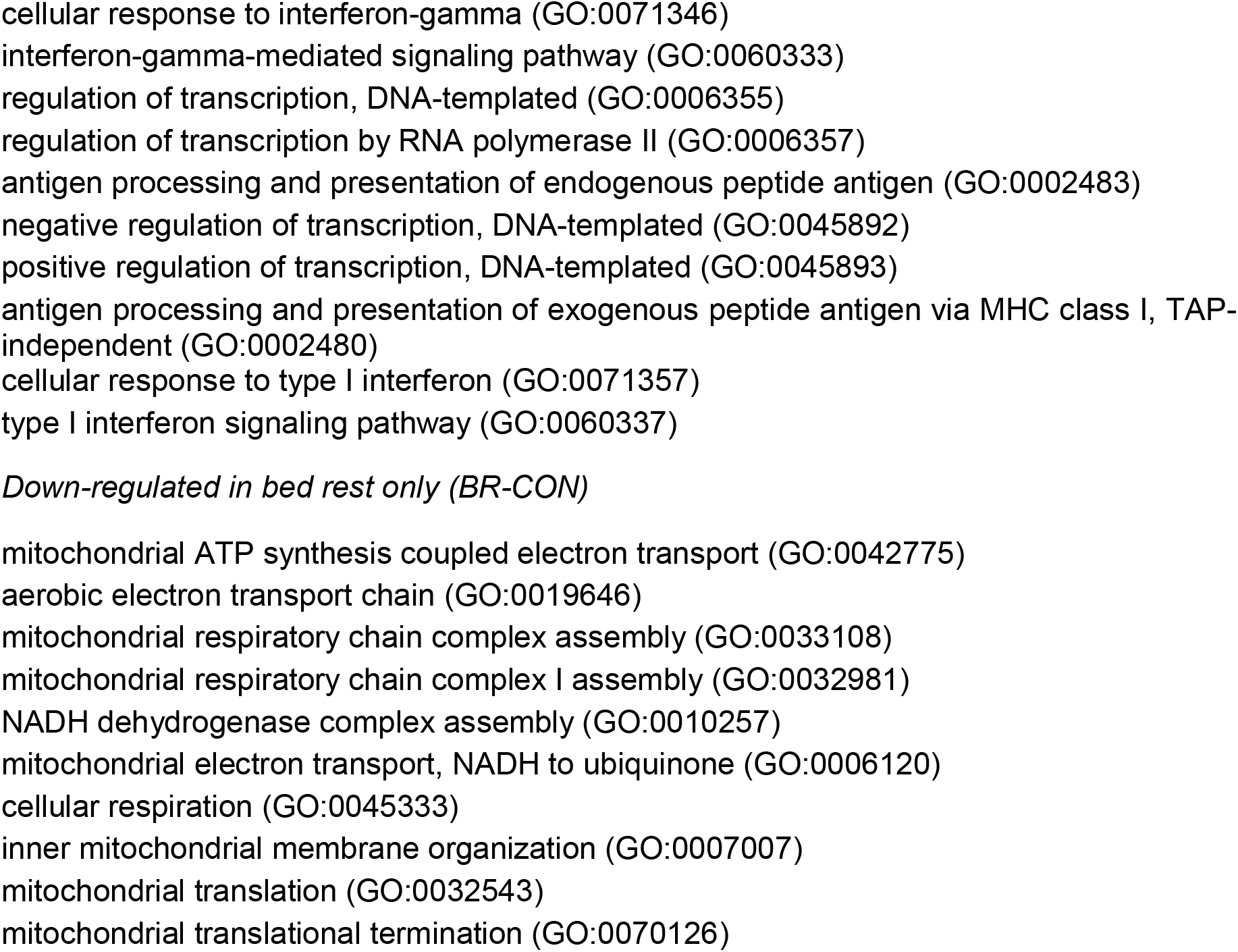
Top GO biological pathways enriched by 35 days of bed rest in BR-CON subjects.

**Table 4:**
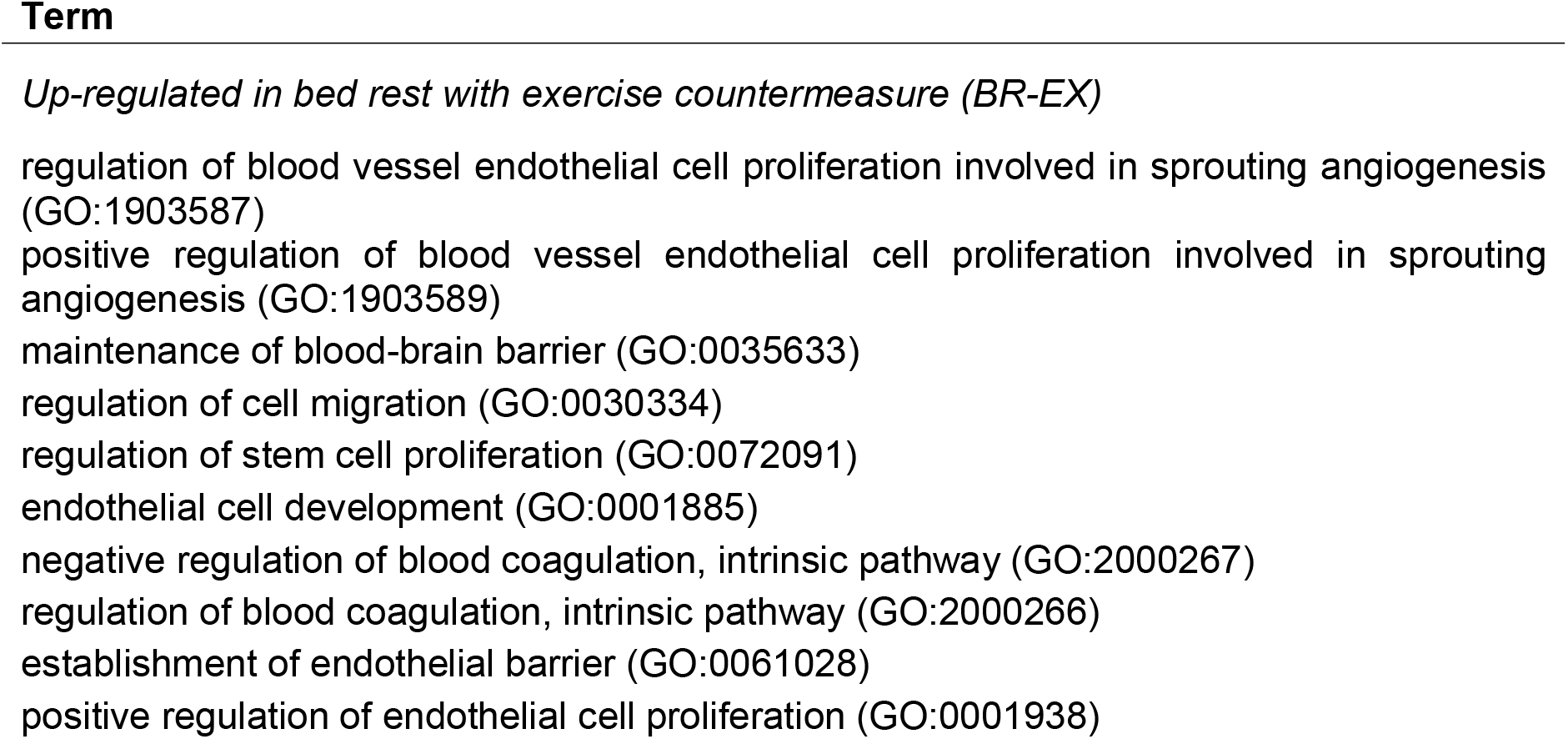
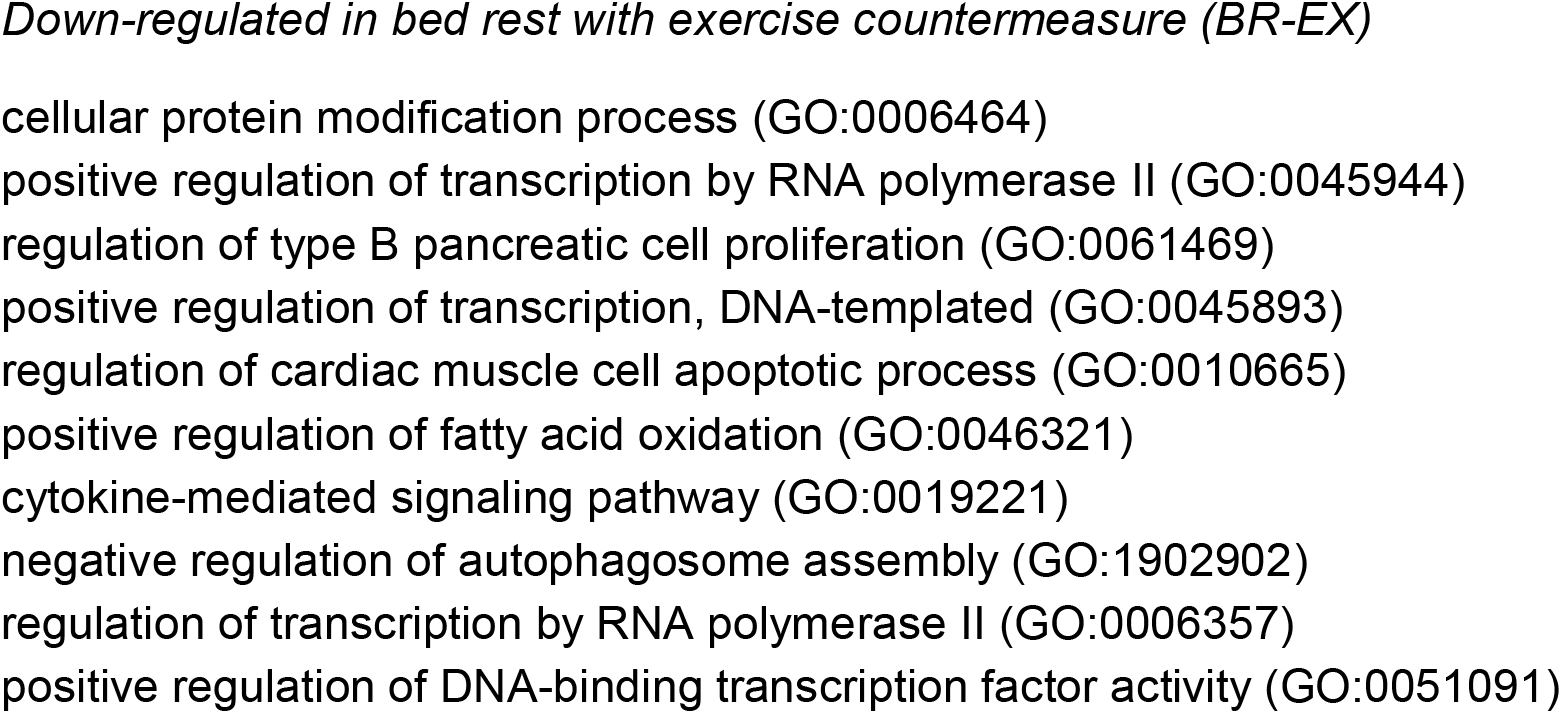
Top GO biological pathways enriched by 35 days of bed rest with an exercise countermeasure in BR-EX subjects.

### Ligand-Receptor Interactome between bed rest DEGs and human dorsal root ganglia neurons

To assess the potential impact of transcriptional changes in muscle from bed rest subjects on the peripheral nervous system, we used a genome-wide ligand-receptor pair database curated for pharmacological interactions relevant to neuro-immune systems (Wangzhou et al., 2021). This interactome permits analysis of a given gene list, and identifies those genes which are capable of ligand-receptor interactions because the ligands have at least one known receptor expressed by human dorsal root ganglia neurons. Up-regulated DEGs from BR-CON and BR-EX were analyzed via the interactome. Of the 591 genes up-regulated post-intervention in BR-CON, 26 genes were found to have known receptors in human DRG (Table 5), including *PGF, IL18, CCL2, EFNA1, CCN1*, and *CXCL2*. In the BR-EX group, there were 7 up-regulated genes post-intervention with known DRG receptors (Table 6).

**Table 5:** Interactome summary of up-regulated BR-CON ligands to human dorsal root ganglia receptors.

| BR-CON up-regulated ligand | DESeq2 adjusted p value | Known DRG receptor(s) |
| --- | --- | --- |
| *ADM* | 8.418E-05 | *RAMP2*, *CALCRL* |
| *ANGPTL2* | 1.452E-03 | *TIE1* |
| *ANGPTL4* | 3.569E-02 | *TIE1* |
| *APOD* | 2.332E-02 | *LEPR* |
| *B2M* | 2.990E-04 | *HLA-F, TFRC, LILRB2, CD3D, LILRB1, CD247, HFE, CD3G, KLRD1, KLRC1* |
| *CALM1* | 2.502E-03 | *PTPRA, TRPV1, SCN10A, GRM4, KCNQ1, GRM7, FAS, PDE1C, PPAPDC2, INSR, PDE1A, PDE1B, EGFR, TRPC3, KCNN4, SELL, KCNQ5, ABCA1, KCNQ3, MYLK, CACNA1C, VIPR1, CRHR1, SCN4A, ADCY8, ADCYAP1R1, HMMR, OPRM1, AQP6, MYLK2, MIP* |
| *CCL2* | 2.725E-03 | *DARC, CCR10, CCR1, CCR2, CCR5* |
| *CCN1* | 4.402E-03 | *ITGB2, CAV1, ITGB5, ITGAV, ITGA5, ITGAM, ITGB3* |
| *CDH15* | 4.944E-02 | *ARVCF, SRPK3* |
| *CX3CL1* | 2.428E-02 | *CX3CR1* |
| *CXCL16* | 1.383E-02 | *CXCR6* |
| *CXCL2* | 4.427E-02 | *CXCR2* |
| *CXCL9* | 7.099E-08 | *CXCR3* |
| *EFNA1* | 1.843E-02 | *EPHB6, EPHA8, EPHA2, EPHA5, EPHA6, EPHB1, EPHA3, EPHA4, EPHA1, EPHA7* |
| *FARP2* | 4.197E-03 | *PLXNA3, PLXNA2, PLXNA1, PLXNA4* |
| *HLA-A* | 1.071E-02 | *APLP2, ERBB2, LILRB2, CD3D, LILRB1, CD3G* |
| *HLA-B* | 4.012E-04 | *CANX, LILRB2, CD3D, LILRB1, CD3G, KLRD1* |
| *HLA-C* | 6.795E-04 | *NOTCH4, DDR1, LILRB2, CD3D, LILRB1, CD3G* |
| *HLA-E* | 1.010E-02 | *SLC16A4, KLRD1, KLRC1* |
| *ICAM2* | 6.004E-03 | *ITGB2, ITGAM, ITGAL* |
| *IL18* | 3.165E-03 | *CD48, IL18R1, IL1RL2, IL18RAP* |
| *LTF* | 4.986E-02 | *LRP1, LRP11, TFRC, GPR162* |
| *PGF* | 2.408E-02 | *NRP2, NRP1, FLT1* |
| *TYMP* | 1.958E-02 | *UPP1, TK2, NT5M, UMPS, TK1, UPRT, DPYD, CDA* |
| *VEGFD* | 3.021E-04 | *KDR, ITGA9, FLT4* |
| *WNT4* | 4.884E-02 | *FZD2, FZD6* |

**Table 6:** Interactome summary of up-regulated BR-EX ligands to human dorsal root ganglia receptors.

| **BR-EX up-regulated ligand** | **DESeq2 adjusted p value** | **Known DRG receptor(s)** |
| --- | --- | --- |
| *A2M* | 4.903E-02 | *LRP1* |
| *ACE* | 3.833E-03 | *BDKRB2* |
| *CXCL12* | 6.281E-03 | *ITGB1, SDC4, CD4, CXCR4, CXCR3* |
| *DLL4* | 2.937E-02 | *NOTCH4, NOTCH1, NOTCH2, NOTCH3* |
| *F8* | 7.898E-03 | *LRP1, LDLR, ASGR2* |
| *JAM2* | 1.585E-03 | *ITGB1, ITGB2, JAM3, TJP1, ITGAM, F11R, ITGA4* |
| *PCDH12* | 7.898E-03 | *PCDH17* |

## Discussion

The analysis described above yields several key conclusions. Muscle biopsy RNA sequencing shows that 5 weeks of bedrest causes a large transcriptional change in muscle that is mostly prevented by an exercise intervention. This transcriptional change in sedentary muscle produces a set of ligands in muscle tissue that may interact with receptors on peripheral sensory neurons that provide sensory and nociceptive information from muscle and/or other tissues that could be exposed to factors released from muscle. This muscle to DRG neuron signaling may play an underappreciated role in the physiological sequalae that emerge from muscle deconditioning (Figure 2).

**Figure 2:**
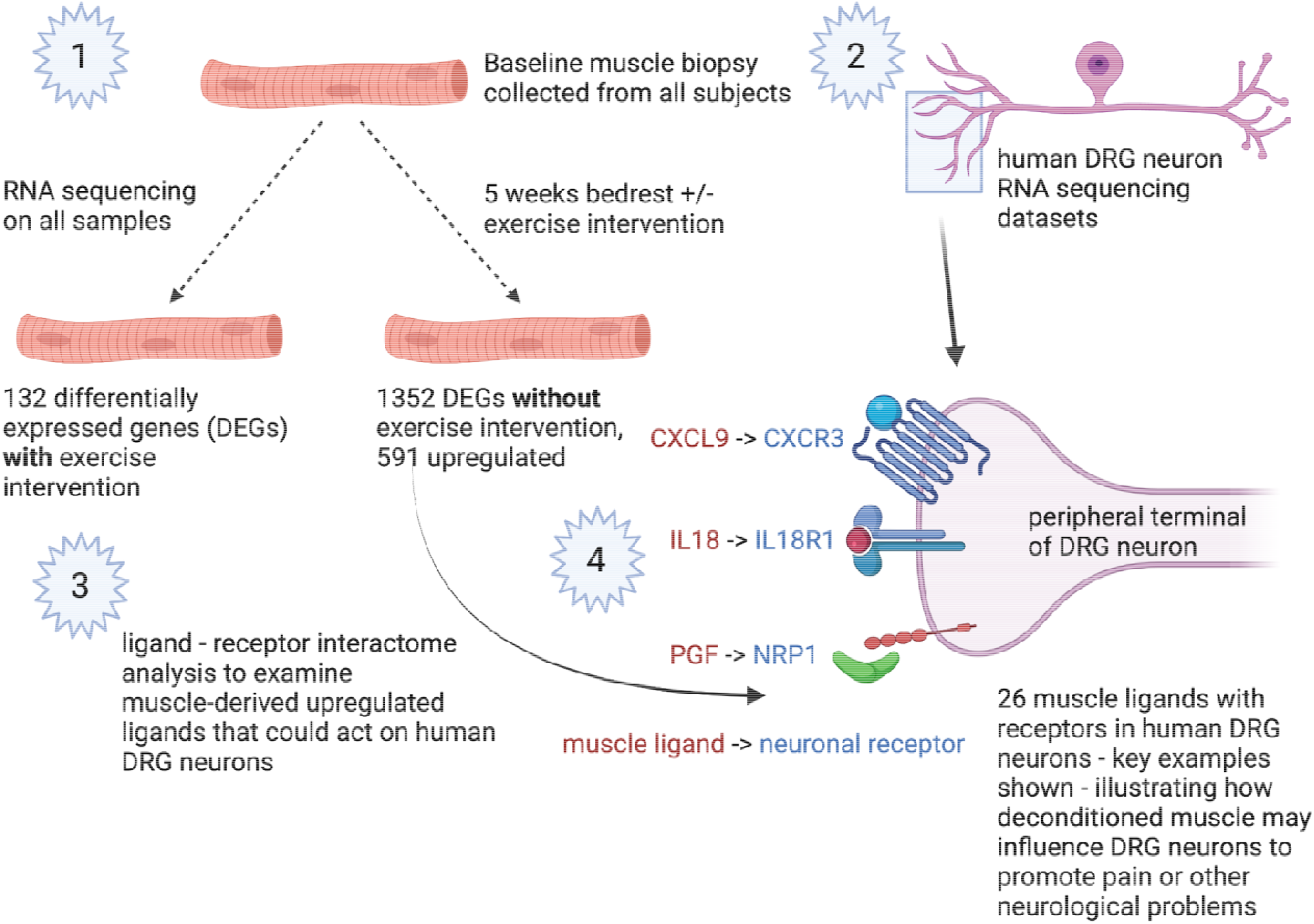
Summary of main findings. (1) Experimental design and top-level findings from RNA sequencing analysis. (2) Pre-existing human DRG sequencing data and a ligand-receptor interactome (3) enabled looking at ligand-receptor interactions between deconditioned muscle and DRG neurons. (4) Key muscle ligands with increased expression in deconditioned muscle (in red) matched to receptors that are expressed by human DRG nociceptors (in blue). Made with Biorender.com.

The findings presented here provide a new perspective on the transcriptional response to bedrest and how this is altered by an exercise intervention. Consistent with other biochemical and physiological measures, exercise during bedrest minimized changes in the muscle transcriptional response, in particular changes that are likely involved in inflammation and signaling to the peripheral nervous system. Many of the genes that were upregulated in muscle samples without exercise intervention have been reported in previous studies, but this study used a longer bedrest interval, and RNA sequencing provides a more comprehensive picture of how gene expression is altered over this time period. A shortcoming is the use of bulk RNA sequencing rather than a single cell approach that could provide more insight into cellular phenotype (Stark et al., 2019). We decided on a bulk RNA sequencing approach for several reasons. First, the samples were collected years prior and stored at −80C so cellular integrity may have been compromised in samples. Second, protocols for nuclei isolation are not optimized for samples that have been frozen for years. Finally, we reasoned that a bulk sequencing approach would give us a fuller picture of transcriptional changes that could be informative for the design of future prospective studies.

We focused our ligand-receptor interaction on muscle to sensory neuron communication because of the link between exercise and pain. A consistent observation in clinical and preclinical pain studies is that exercise is one of the most successful interventions for chronic pain (Sluka et al., 2018; Lesnak and Sluka, 2020; Merkle et al., 2020). One potential explanation for this effect is the production of myokines from exercise that act on sensory neurons to decrease their excitability (Sharma et al., 2010). Another is that lack of exercise, the primary cause of muscle deconditioning, could produce factors from muscle tissue that promote pain through an action on DRG sensory neurons. Our hypothesis generating analysis provides a framework for testing this latter idea in mechanistic studies. We found a number of ligands, including chemokines (*CXCL2* and *CXCL9*), interleukins (*IL18*) and growth factors (*PGF*) that were strongly upregulated in the BR-CON group that have cognate receptors that are expressed by human sensory neurons, including sensory neurons called nociceptors that play a key role in detecting pain and are sensitized in chronic pain states (Figure 2). IL-18 has been associated with pain promotion in animal models (Verri et al., 2007) and in cancer pain patients (Heitzer et al., 2012). IL-18 receptors are expressed in human DRG but IL-18 may also act indirectly through endothelin receptors (Verri et al., 2004), which are robustly expressed by human nociceptors (Tavares-Ferreira et al., 2021). Placental growth factor (PGF) is a vascular endothelial growth factor (VEGF) family growth factor that can act via neuropilin receptor (*NRP1* and *NRP2* genes) complexes (Dewerchin and Carmeliet, 2014). NRP1 was recently shown to play an important role in promotion of neuropathic pain in rats (Moutal et al., 2020) and is also highly expressed by human nociceptors (Tavares-Ferreira et al., 2021). PGF isoforms that encode peptides with heparin binding functionality, and are able to interact with Neuropilin-1 and Neuropilin-2, were found to be robustly detected (Reads per Million mapped reads or RPM > 0.5 in 22 out of 26 samples, Supplementary Data, Sheet 1). Finally, while interferon γ was not directly detected as an upregulated gene, downstream interferon γ signaling was the most enriched GO term detected in the BR-CON group. Muscle deconditioning may promote increased interferon γ expression in other cells types that can then act on human nociceptors which express interferon γ receptors (Tavares-Ferreira et al., 2021). Importantly, interferon γ has been positively associated with several clinical pain conditions (Parkitny et al., 2013; Kamieniak et al., 2020). These ligand-receptor interactome findings lay a foundation for future hypothesis-based work using human DRG neurons (Renthal et al., 2021) to better understand how deconditioned muscle may signal to the peripheral nervous system.

The data and analyses described here are a useful resource for understanding how muscle responds to long-lasting inactivity. As noted in our results, we were able to accurately classify groups based on their transcriptional response, but we incorrectly called the exercise intervention group prior to unblinding. This mistake was made based on our *a priori* assumption that anti-inflammatory myokines would be produced by exercised muscle. Instead, we discovered that deconditioned muscle produces a broad set of secreted molecules, including a mix of inflammatory and anti-inflammatory mediators, which are likely to have a broad impact on many cell types in the body. Therefore, our unbiased approach reveals new insight into the potential impact that deconditioned muscle may have on the physiology of many organ systems. While our analysis has focused on DRG neurons, our data can be mined for additional contexts. Our robust results with a relatively small sample size suggest that future prospective studies can utilize single cell approaches to build substantially on the findings described here.

